# Nuclear Respiratory Factor 1 (NRF-1) Controls the Activity Dependent Transcription of the GABA-A Receptor Beta 1 Subunit Gene in Neurons

**DOI:** 10.1101/321257

**Authors:** Zhuting Li, Meaghan Cogswell, Kathryn Hixson, Amy R. Brooks-Kayal, Shelley J. Russek

## Abstract

While the exact role of β1 subunit-containing GABA-A receptors (GABARs) in brain function is not well understood, altered expression of the β1 subunit gene (*GABRB1*) is associated with neurological and neuropsychiatric disorders. In particular, down-regulation of β1 subunit levels is observed in brains of patients with epilepsy, autism, bipolar disorder, and schizophrenia. A pathophysiological feature of these disease states is imbalance in energy metabolism and mitochondrial dysfunction. The transcription factor, nuclear respiratory factor 1 (NRF-1), has been shown to be a key mediator of genes involved in oxidative phosphorylation and mitochondrial biogenesis. Using a variety of molecular approaches (including mobility shift, promoter/reporter assays, and overexpression of dominant negative NRF-1), we now report that NRF-1 regulates transcription of *GABRB1* and that its core promoter contains a conserved canonical NRF-1 element responsible for sequence specific binding and transcriptional activation. Our identification of *GABRB1* as a new target for NRF-1 in neurons suggests that genes coding for inhibitory neurotransmission may be coupled to cellular metabolism. This is especially meaningful as binding of NRF-1 to its element is sensitive to the kind of epigenetic changes that occur in multiple disorders associated with altered brain inhibition.

## INTRODUCTION

The type A γ-aminobutyric acid receptor (GABAR) is a ligand-gated Cl^-^ ion channel that mediates inhibitory neurotransmission in the adult mammalian central nervous system. The majority of GABARs are composed of two α and two β subunits, and either a γ2 or δ subunit (Barrera *et al*. 2008, Patel *et al*. 2014, Farrar *et al*. 1999). For each receptor, there is the binding of two molecules of GABA, one molecule at each α and β subunit interface (Connolly & Wafford 2004, Olsen & Sieghart 2009, Rabow *et al*. 1995). In the mature neuron, activation of GABARs leads to hyperpolarization. Depending on its subunit composition, GABARs may contain binding sites for barbiturates, benzodiazepines, ethanol, and/or neuroactive steroids. There are nineteen different subunit genes to date, grouped into eight classes (i.e. α1–6, β1–3, γ1–3, δ, ε, θ, π, ρ1–3) that contribute to the diversity and differential assembly of receptor subtypes. The β subunits, which contain the domains that interact with mediators of receptor trafficking and endocytosis (for reviews see (Vithlani *et al*. 2011, Jacob *et al*. 2008)), play an important role in the expression of GABARs at the cell surface.

The human *GABRB1* gene, located on chromosome 4, is part of a GABAR gene cluster that contains the genes that encode the α2, α4, and γ1 subunits. A dysregulation of GABAR-mediated neurotransmission has been implicated in various neurological disorders (Hines *et al*. 2012) that show altered levels of GABAR subunits, including β1. Through linkage studies, *GABRB1* has been associated with alcohol dependence (Parsian & Zhang 1999, Sun *et al*. 1999, Zinn-Justin & Abel 1999, Song *et al*. 2003); and more recently, specific mutations in mouse *Gabrb1* have been shown to produce increased alcohol consumption that is linked to increased tonic inhibition (Anstee *et al*. 2013). Interestingly, single nucleotide polymorphisms in *GABRB1* are also associated with altered brain responses in human adolescents susceptible to addictive behaviors (Duka *et al*. 2017).

*GABRB1* expression is also reduced in the lateral cerebella of subjects with bipolar disorder, major depression, and schizophrenia compared to healthy subjects (Fatemi *et al*. 2013). Particularly in schizophrenia, a significant association of *GABRB1* has been identified by genome-wide association studies that were coupled to a protein-interaction-network-based analysis (Yu *et al*. 2014). As *GABRB1* and *GABRA4* lie within the same GABAR gene cluster and their promoters are head-to-head, it is interesting to note that the association of *GABRA4* with autism risk increases with a *GABRB1* interaction (Ma *et al*. 2005, Collins *et al*. 2006), suggesting that these genes may be coordinately regulated. Further support for an association of *GABRB1* with autism is evidenced by a decrease in β1 subunit levels in the brains of autistic subjects (Fatemi *et al*. 2009, Fatemi *et al*. 2010). In addition, the levels of both β1 and β2 subunit mRNAs are reduced in a Fragile X mental retardation mouse model, where the gene Fragile X mental retardation 1 (fmr1) was removed (D’Hulst *et al*. 2006). Finally, down-regulation of β1 subunit mRNAs and protein are observed in the rat pilocarpine model of epilepsy (Brooks-Kayal *et al*. 1998). Yet, despite its prevalent association with brain disorders, there is still little known about the function and/or regulation of β1 in neurons.

The TATA-less *GABRB1*/*Gabrb1* promoter (*GABRB1-p* (human)*/Gabrb1-p* (rodent)) contains multiple transcriptional start sites that lie within a CpG island (Russek *et al*. 2000, Saha *et al*. 2013). In unraveling the molecular determinants of GABAR β1 subunit gene regulation, our laboratory demonstrated that the minimal *GABRB1-p* lies within the first 500 bp of the 5’ flanking region. Within this region, there is a conserved initiator element (Inr) that mediates down-regulation in response to chronic GABA exposure, implicating an autologous mechanism of transcriptional control.

Nuclear respiratory factor 1 (NRF-1) is a transcription factor that functions primarily as a positive regulator of nuclear genes involved in mitochondrial biogenesis and oxidative phosphorylation, such as Tfam, which moves into the mitochondria and regulates mitochondrial DNA transcription (Scarpulla 2006, Scarpulla 2008). However, it has also been shown that the binding of NRF-1 to a co-factor, such as SIRT7, can influence its polarity (from activator to repressor) (Mohrin et al., 2015). In addition, binding of NRF-1 to DNA is regulated by the methylation state of its regulatory element (Domcke et al., 2015), suggesting that its role in neuronal gene expression will be sensitive to the epigenetic changes that occur in neurological and neuropsychiatric disorders.

It is well known that increased neuronal activity results in a parallel change in cellular metabolism, as orchestrated by the synthesis of NRF-1 and its control over mitochondrial biogenesis. Moreover, it has been reported that NRF-1 is a transcriptional activator of glutamate receptor subunit genes under conditions of depolarizing stimulation in neurons (Dhar & Wong-Riley 2009) suggesting that in addition to its role in cellular metabolism, via regulation of the mitochondrial genome, NRF-1 coordinates activities in the nucleus to couple neuronal excitability with energy demands of synaptic neurotransmission.

Here, we ask whether NRF-1 may control the transcription of GABAR subunit genes (*GABRs*), and in particular the human β1 subunit gene (*GABRB1*), a gene that has been associated with neuronal developmental disorders, the pathophysiology of epilepsy, and alcohol dependence. In this study, we have uncovered a functional regulatory element within *GABRB1* that demonstrates sequence specificity and is responsible for the majority of *GABRB1* promoter-reporter activity, as well as a role for NRF-1 in the activity dependent transcription of endogenous *Gabrb1* in rat primary cortical neurons.

## METHODS

*Cell Culture and Drug treatment—* <u>The use of animals for our culture studies was under the guidance and protocol approval of the Boston University Institutional Care and Use Committee (IACUC).</u>

Primary neocortical neurons were isolated from embryonic day 18 Sprague-Dawley rat embryos (Charles River Laboratories). Isolated embryonic brains and the subsequently dissected cortices were maintained in ice-cold modified calcium-magnesium free Hank’s Balanced Salt Solution (HBSS) (4.2mM sodium bicarbonate, 1 mM sodium pyruvate, and 20 mM HEPES, 3mg/ml BSA) buffering between pH range 7.25–7.3. Tissues were then separated from HBSS dissection solution and trypsinized (0.05% trypsin-EDTA) for 10 minutes in 37^°^C and 5% CO_2_. The trypsin reaction was stopped with serum inactivation using plating medium (Neural Basal Medium, 10% FBS, 10 U/ml penicillin/streptomycin, 2mM L-glutamine). Tissues were triturated with a 1000 mL micropipette and diluted to a concentration of 0.5x10^6^ cells/mL in plating media for plating. Cells were allowed to adhere onto Poly-L-lysine coated culturing surface for 1 h prior to changing to serum-free feeding medium (2% B-27, 2mM glutamine, 10 U/ml penicillin/streptomycin supplemented neurobasal medium). Neuronal cultures were maintained at 37^°^C in a 5% CO_2_ incubator. Primary cortical neurons (DIV7-8) were treated with either Vehicle or 20 mM KCl for 6 hours before harvesting for analysis.

*Expression Constructs—* pCDNA 3.1 hygro hNRF-1 VP16 was generously provided by Dr. Tod Gulick (Ramachandran *et al*. 2008) (Sanford-Burnham Medical Research Institute, Orlando FL). pcDNA 3.1 hygro hNRF-1 VP16 encodes a constitutively active form of NRF-1, consisting of the full-length human NRF-1 and the herpes simplex virus VP16 transactivation domain. The pcDNA3.0-NRF-1 DN expresses amino acid residues 1-304 of human NRF-1, which encodes the DNA-binding, dimerization and nuclear localization domains of NRF-1, but lacks the transactivation domain (amino acids 305-503). With the exception of a single conservative mutation at amino acid residue 293 (A→T), the 304 amino acid residues of NRF-1 are conserved between human and rat. The construct was created using PCR with the forward primer sequence 5’-CGG<u>GGTACC</u>ACCATGGAGGAACACGGAGTGACCCAAAC-3’, containing the underlined Kpn1 restriction site and the kozak sequence on the 5’ end, and the reverse primer 5’GC<u>TCTAGA</u>TCACTGTGATGGTACAAGATGAGCTATACTATGTGTGGCTGTGGC-3’, containing stop codon and Xba1 restriction site. PCR products were digested with restriction enzymes Kpn1 and Xba1, and ligated into pcDNA3.0 vector (Invitrogen).

*Electrophoretic mobility shift (EMSA) and supershift assays*—Briefly, 30 bp DNA probes containing the putative NRF-1 binding sequence were incubated with 25µg of neocortical nuclear extracts for electrophoresis under non-denaturing conditions. Following electrophoresis, the protein-DNA complexes were detected by autoradiography. The DNA probes were created from annealing synthesized oligonucleotides (www.idtdna.com) and 5’ end labeling using [γ-³²P] ATP (PerkinElmer) in a T4 polynucleotide kinase (NEB) reaction. Nuclear extracts were prepared from DIV7 primary neocortical neurons grown on 10-cm plates in the presence of protease inhibitor cocktail. Protein-DNA binding specificity was determined by adding poly (dI-dC) (Roche) or/and 100-fold excess unlabeled DNA probe prior to the addition of labeled probe during the room temperature binding reaction. To generate a supershift complex, NRF-1 antibody (AbCam ab34682) was added to the reaction mixture for 15 min. The binding reactions were loaded onto a 5% polyacrylamide gel in 0.5X TBE buffer and run at 200V for 2 h at 4^°^C. The positive control probe consisted of a functional NRF-1 sequence (Evans & Scarpulla 1990) found in the Rat cytochrome c (rCycs) gene (Evans & Scarpulla 1990). Probe and competitor oligonucleotide sequences were: *GABRB1* NRF1, 5’-agcgcgc**TCTGCGCATGCGCA**ggtccattc-3’ and 5’-gaatggacc**TGCGCATGCGCAGA**gcgcgct-3’. *GABRB1* NRF1 mutant, 5’-agcgcgc**TCTGC**c**CATG**g**GCA**ggtccattc –3’ and 5’-gaatggacc**TGC**c**CATG**g**GCAGA**gcgcgct-3’. *rCycs* NRF1, 5’-ctgcta**GCCCGCATGCGC**gcgcacctta-3’and 5’-taaggtgcgc**GCGCATGCGGGC**tagcag-3’.

*Reporter Plasmids and Promoter Mutagenesis*—The *GABRB1p*-Luc (pGL2-*GABRB1*) promoter construct containing the 5’ flanking region of the human β1 subunit gene was previously cloned by our laboratory and contains 436 bp upstream of the initiator sequence and 105 bp downstream (Russek et al.

2000). The promoter containing a mutated NRF-1 element (**TCTGC**c**CATG**g**GCA**) within the *GABRB1p-Luc* was created by PCR-driven overlap extension. Using wild-type *GABRB1p*-Luc as PCR template, two PCR fragments were amplified using the GL1 primer (Promega) and the antisense mutant NRF-1 oligonucleotide from EMSA, and sense mutant NRF-1 oligonucleotide and the GL2 primer (Promega), resulting in fragments with 30 bp overlapping sequences that contain the mutant NRF-1 element. A second PCR step using GL1, GL2 primers and both initial PCR products produced the mutant *GABRB1* promoter insert.

*Luciferase Assay/Reporter Assay with Magnetofection*™— Magnetofection of DNA into primary neuron cultures was achieved with the NeuroMag transfection reagent according to the manufacturer’s protocol. Here, 2 ml of resuspended E18 primary cortical neurons at 0.5 X 10^6^ cells/ml were plated in each well of a 6-well plate. On DIV7, neurons were transfected with 1µg of expression construct, 2µg of promoter reporter construct, and 3µl of NeuroMag transfection reagent (1:1 DNA to reagent ratio). 24 h after transfection, neurons were actively lysed by scraping. Cell lysates were cleared of precipitates by centrifugation and then assayed for luciferase activity using a luciferase assay system (Promega). Luciferase activity was normalized to total protein as determined using a protein assay kit (Thermo Scientific Pierce). All transfections were performed in sister dishes from three or more plating sessions to produce true N’s.

*Chromatin Immunoprecipitation (ChIP)—* ChIP was performed according to the Magna ChIP A protocol (Millipore). Briefly, primary neurons in 100 mm dishes were fixed with a final concentration of 1% formaldehyde in culturing media. The remaining unreacted formaldehyde was quenched with Glycine. Genomic DNA and protein complexes were extracted from cells using nuclear lysis buffers supplemented with protease and phosphatase inhibitors. The lysates containing DNA-protein complexes were sonicated (nine times, 5 minutes each at a 30s on/off interval) in an ice-cold water bath with a Bioruptor (Diagenode) in order to generate fragments predominantly in the range of 200–500 bp in size. The sheared chromatin was immunoprecipitated with either anti-NRF-1 antibody (Abcam ab34682 ChIP grade antibody) or normal rabbit IgG overnight at 4ΰC with constant rotation. The antibody/transcription factor bound chromatin was separated from unbound chromatin using Protein A conjugated magnetic beads and magnetic pull-down. The isolated complexes were washed with a series of salt buffer solutions prior to eluting. DNA fragments were separated from complexes using Proteinase K and heating, and recovered through column purification. The co-precipitated DNA fragments were identified by quantitative PCR (qPCR) using specific primers and TaqMan probes that flank putative responsive elements in gene promoters using the FastStart Universal Probe Master (Roche) PCR reagent. PCR cycling was performed using the ABI7900HT Fast Real-Time PCR system. The *Gabrb1* promoter fragment (114 bp) was amplified using: forward primer 5’- TGTTTGCAAGGCACAAGGTGTC-3’, reverse primer 5’- TCTGCGAAGATTCAAGGAATGCAACT, TaqMan^®^ MGB probe 5’- GCGCATGCGCAGGTCCATTCGGGAAT-3’.

*Western Blot Analysis*—Total cellular proteins were extracted from primary neuronal cultures after KCl treatment with standard procedures and the use of RIPA lysis buffer (Tris, pH 7.4, 10 mM; Nonidet P-40 1%; NaCl 150 mM; SDS 0.1%; protease inhibitor mixture (Roche Applied Science) 1X; EDTA 1mM; sodium orthovanadate 1 mM; sodium deoxycholate 0.1%; phenylmethylsulfonyl fluoride 1 mM). 30 µg of whole cell extracts were separated by SDS-PAGE under reducing conditions on either 10% or 4-20% Tris-glycine gel according to mass/size. The electrophoresed samples were transferred to nitrocellulose membranes. Western blot analysis was performed using antibodies against NRF-1 (AbCam ab34682, 1:2000 in 1X TBS-T). Membranes were incubated with peroxidase-conjugated goat anti-rabbit secondary antibody (Santa Cruz Biotechnology, 1:5000) in TBS-T and visualized using the ECL enhanced chemiluminescence reagent (GE Healthcare Life Sciences). Data are presented as mean ± SEM. Significance was set at p < 0.05, as determined using the paired Student’s t-test (two-tailed).

*RNA extraction and qRT-PCR—*Total RNA was isolated from cultured primary neocortical using the RNeasy Micro Kit (Qiagen). For each reaction, 20ng of total RNA was reverse-transcribed to cDNA and PCR amplified in a single reaction mixture using the TaqMan^®^ One-Step RT-PCR Master Mix Reagents Kit (Applied Biosystems). Incubation and thermal cycling conditions were performed using the ABI7900HT in a 384-well PCR plate format (AppliedBiosystems). The RT reaction was held at 48^°^C for 30 min, followed by 95^°^C for 10 min to activate the polymerase. The PCR reaction conditions were: 15 sec denaturation at 95^°^C and coupled annealing and extension for 1 min at 60^°^C for 40 cycles. Co-detection of rat peptidylprolyl isomerase A (cyclophilin A) gene served as an internal control for normalization. Cyclophilin A expression has been shown to be stable in response to neuronal stimulation in culture (Santos & Duarte, 2008), which is consistent with our previous studies. Relative gene expression was quantified using 2^ (-ΔΔC_T_) and a standard curve was generated based on the amplification of total RNA extracted from untreated cultured neurons. The qRT-PCR primers and probes for rat mRNAs were: *NRF-1*, 57 bp amplicon (Assay ID: Rn01455958_m1, Thermo Fisher Scientific); *Gabrb1*, 81 bp amplicon (Assay ID: Rn00564146_m1, Thermo Fisher Scientific); *Ppia*, 60 bp amplicon, forward primer: 5’- TGCAGACATGGTCAACCCC-3’, reverse primer: 5’- CCCAAGGGCTCGCCA-3’, TaqMan probe with TAMARA quencher: 5’- CCGTGTTCTTCGACATCACGGCTG-3’.

## RESULTS

### Neuronal depolarization increases NRF-1 and GABAR β subunit gene transcription

To determine whether *Gabrb1* is activity dependent, primary cortical neurons were treated with KCl. Both NRF-1 protein and NRF-1 mRNA levels have been previously shown to increase with KCl-stimulated depolarization (Dhar & Wong-Riley 2009). We asked whether under conditions where NRF-1 levels increase in response to neuronal activity, is it accompanied by increased levels of *Gabrb1* transcripts. As shown in Figure 1A and 1B, there is a 2-fold increase in NRF-1 mRNA levels (1.982 ± 0.445, n=5, **p < 0.01) upon KCl stimulation for 6 hours that is accompanied by a ∽30% increase in the levels of NRF-1 protein (fold change: 1.285 ± 0.330, n=6, *p < 0.05) when compared to vehicle control. In parallel to changes in NRF-1, we now report a ∽40% increase in levels of *Gabrb1* transcripts (fold change: 1.424 ± 0.324, n=5, *p < 0.05).

**Figure 1.**
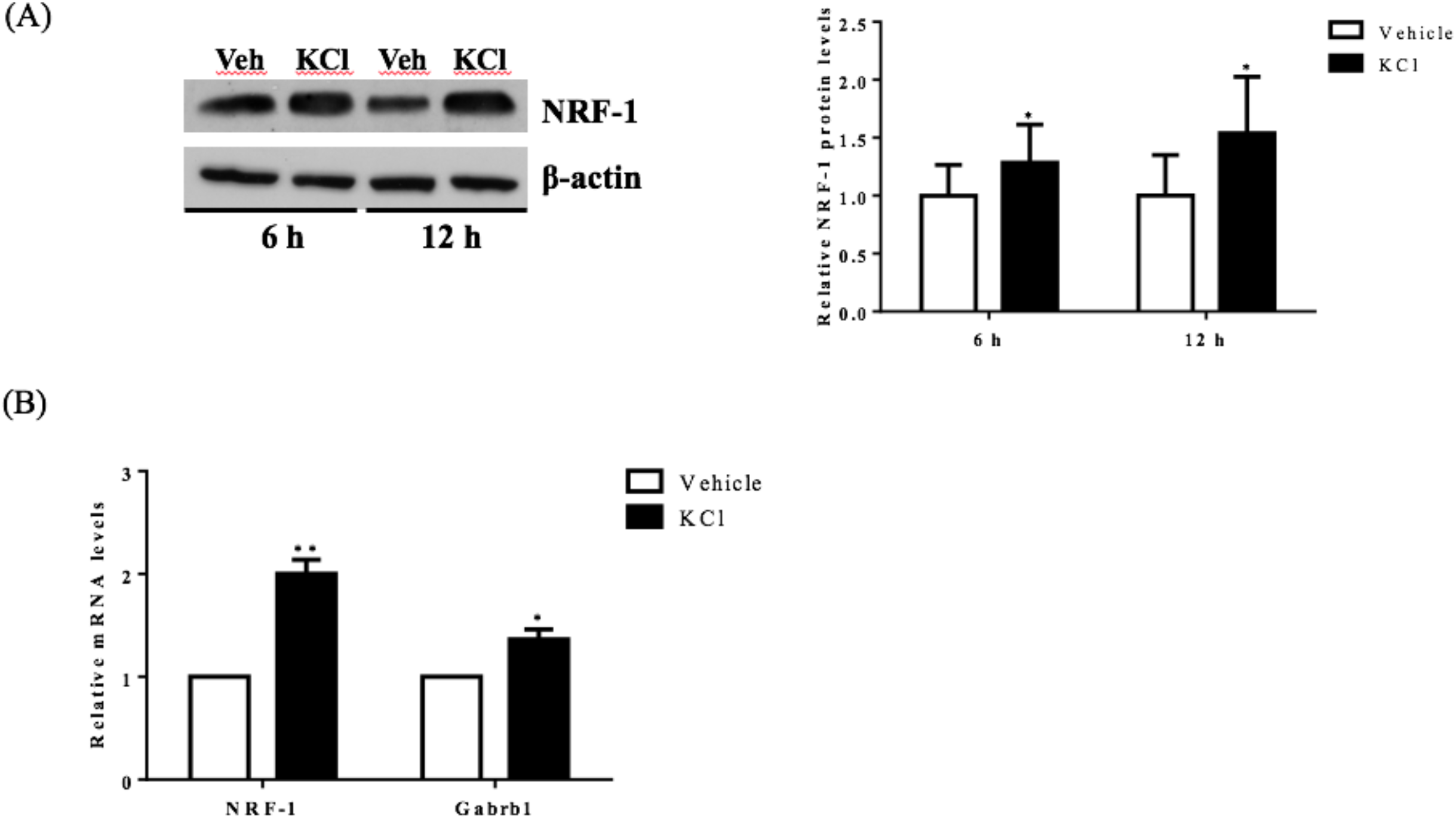
Activity-dependent regulation of *NRF-1* and *Gabrb1* in primary cortical neurons. Primary cortical neurons (DIV7–8) were treated with either Vehicle or 20 mM KCl for 6 hours. (A) Total protein was extracted from neurons and probed for the presence of NRF-1 and β-actin. A representative western blot is shown (*left*) for comparison. NRF-1 levels were quantified by densitometry and normalized to levels of β-actin. Levels of NRF-1 are expressed relative to vehicle (*right*) (n = 6, paired Students t-test, p<.05 as significant. (B) Levels of mRNAs were quantified by TaqMan qRT-PCR. Transcripts specific to *NRF-1* and *Gabrb1* were normalized to Cyclophilin A. Messenger RNA levels are expressed relative to vehicle treated neurons plated in a 6 well dish. Data represent the average ± SEM of n = 6 independent neuronal cultures with neurons extracted from different animals and plated on different days. *, p < 0.05, **, p < 0.01, Student’s t-test. This dataset was originally published in the thesis of Li, Zhuting (2015) Activity-dependent gene regulation in neurons: energy coupling and a novel biosensor [dissertation]. [Boston (MA)]: Boston University.

### Identification of a Conserved NRF1 Element in the GABRB1 Promoter

Our laboratory previously defined the 5’-regulatory region of the human β1 subunit gene *GABRB1*, identifying transcriptional start sites (TSSs) within a 10 bp functional initiator element (Inr) that mediates the response of the gene to chronic GABA exposure (Russek et al. 2000, Saha et al. 2013). Now we report that directly upstream of this Inr is a canonical NRF-1 element spanning −11/+1 relative to the major TSS for the rat homologue *Gabrb1* in neocortical neurons. As shown in Figure 2, the location of the NRF-1 element within the promoter region is conserved across multiple species. Given the ubiquitous expression of NRF-1, its conservation across species, and its established role in cellular respiration and mitochondrial biogenesis, the sequence comparison presented in Figure 2 strongly suggests that the NRF-1 element is functionally relevant to β1 subunit expression in the mammalian brain.

**Figure 2.**
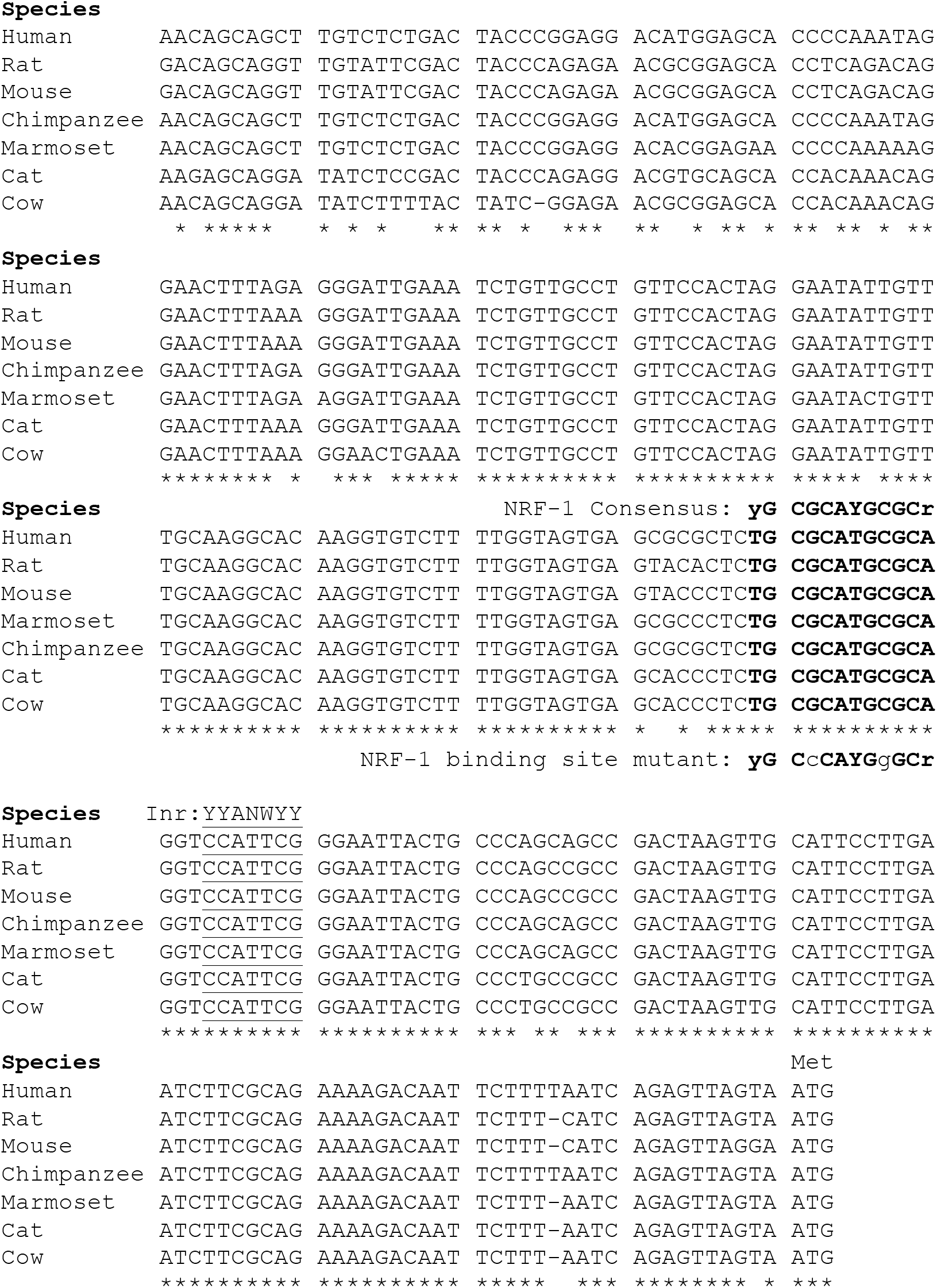
Sequence alignment of the 5’ promoter regions of β1 subunit genes. The β1 subunit promoters in mammals contain a conserved NRF-1 element, indicated in bold type upstream of the major initiator element (Inr) specific to each gene, underlined for reference. Sequences were aligned using ClustalW, where conserved nucleotides are as indicated “*”. Modified from the figure originally published in the thesis of Li, Zhuting (2015) Activity-dependent gene regulation in neurons: energy coupling and a novel biosensor [dissertation]. [Boston (MA)]: Boston University.

### NRF-1 Recognizes the Cis-Element in the human GABRB1 Promoter

To determine the specific binding site within *GABRB1-p* that binds to NRF-1, we performed an electrophoretic mobility shift assay (EMSA) with a ^32^P-labeled probe specific to its NRF-1 consensus element in a binding reaction with nuclear protein extracts from E18 primary cortical neurons. To validate the specificity of the NRF-1 antibody for EMSA analysis, nuclear extracts were incubated with a positive control probe (Dhar *et al*. 2008) containing the NRF-1 binding site of the rat cytochrome c promoter. As shown in lane 2 of Figure 3A, the control radiolabeled probe (rat Cyt C) displays specific DNA recognition from nuclear extracts of cortical neurons that is confirmed by supershift with the addition of an NRF-1 specific antibody (Fig. 3A, lane 4). Next, specific binding to the putative NRF-1 consensus site in *GABRB1* was confirmed using the same nuclear extracts, with sequence specificity defined by competition with an unlabeled double stranded oligonucleotide that was identical to the probe sequence (competitor) (Fig. 3A, lanes 6 and 7). Addition of an unlabeled competitor mutant probe, containing substitutions within the GC core, failed to compete for complex formation (Fig. 3A, lane 8). Presence of endogenous NRF-1 at the *GABRB1* NRF1 consensus site was further confirmed by supershift analysis using the NRF-1 specific antibody (Fig. 3A, lane 9). Finally, a radiolabeled probe containing the sequence of the mutant NRF-1 site in *GABRB1* shows little or no complex formation (Fig. 3A, lanes 10–12).

**Figure 3.**
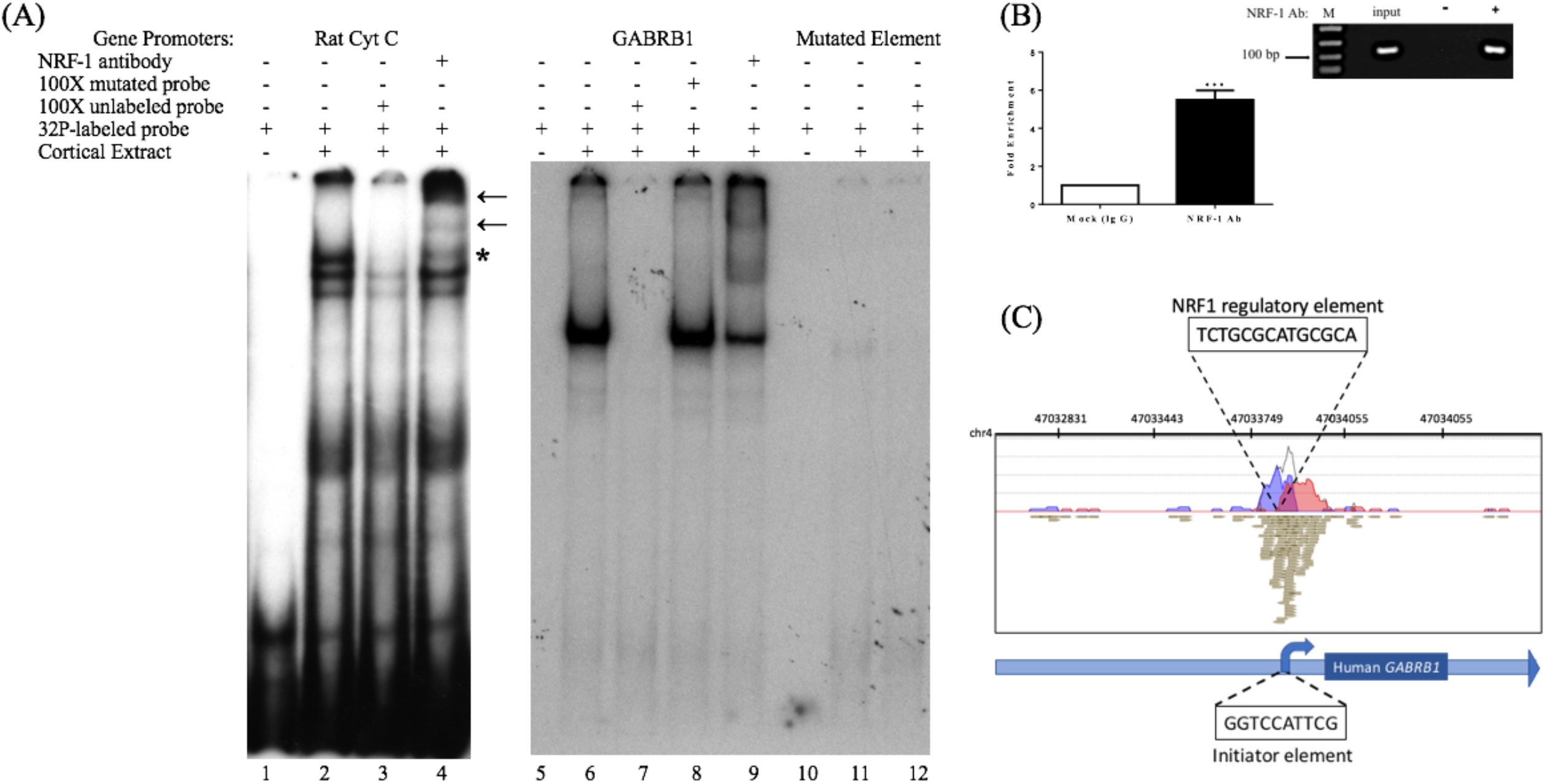
*In vitro and in vivo* binding of NRF-1 to the putative NRF-1 site in *GABRB1*. (A)^32^P-labeled probes encompassing the NRF-1 binding site were incubated with 20 µg of DIV7 primary rat cortical nuclear extracts. 100-fold excess of unlabeled probe was added to the binding reaction to assess specificity. NRF-1 Abs were pre-incubated with nuclear extracts and radiolabeled probe to test for “supershift” and protein identification. (Left Panel) The NRF-1 element in the rat cytochrome c (Cyt C) promoter displays NRF-1 specific binding (lane 2) and “supershift” (lane 4). (Right Panel) The proposed NRF-1 element in the human *GABRB1* promoter displays a probe specific shift (lane 6) (note that excess probe was run off of the gel to provide room for the detection of the shifted probe), competition of complex formation with cold competitor (lane 7), lack of competition with mutant cold competitor (lane 8), and supershift upon addition of NRF-1 specific Ab (lane 9). In contrast, binding to radiolabeled probe for NRF-1 mutant *GABRB1* shows markedly reduced signal strength (lanes 11 and 12). “*” indicates specific interaction between labeled probe and nuclear extract, “←” indicates location of supershift. (B) Chromatin Immunoprecipitation (ChIP) assays were performed using sonicated genomic DNA from DIV7 primary rat cortical neurons and either ChIP grade NRF-1 polyclonal antibody (Abcam, ab34682) that recognizes the full length protein or rabbit IgG (Vector Laboratories, I-1000). Co-precipitated *Gabrb1* gene promoter fragments were detected with specific qPCR primers and probe. Data represent the average ± SEM of n = 4 independent primary cultures and co-precipitations. *, p < 0.05, student t-test. (C) Representative ChIP-seq track from the Strand NGS software platform for *GABRB1* in H1-hESC cells after peak detection (MACS version 2.0). Read density profile plots of forward reads (blue) and reverse reads (red) aligned to the UCSC transcript model are depicted; each brown box represents a single 27-bp sequencing read. The NRF-1 motif sequence is shown in black text above its position. Relative position of the Inr in *GABRB1* is shown for reference. Chr: chromosome. A and B datasets were originally published in the thesis of Li, Zhuting (2015) Activity-dependent gene regulation in neurons: energy coupling and a novel biosensor [dissertation]. [Boston (MA)]: Boston University.

To determine whether the endogenous β1 promoter in neurons is occupied by NRF-1, ChIP was performed using genomic DNA derived from E18 rat primary cortical cultures (DIV7) that was precipitated with NRF-1 antibodies. Precipitated fragments were detected using PCR primers that specifically amplify DNA encompassing the putative NRF-1 binding site in rat *Gabrb1*. As can be seen in Figure 3B, there is a 5-fold increase (5.045 ± 0.981, n=4, *p < 0.05) in PCR detection of the NRF-1 site in *Gabrb1* when precipitated using an NRF-1 Ab, as compared to rabbit IgG. Moreover, NRF-1 is also present at the core promoter of *GABRB1* in human embryonic stem cells (H1-hESC) as detected in ChIP-sequencing (ChIP-seq) datasets of the ENCODE project (https://www.encodeproject.org) using our bioinformatic analysis algorithm (in the Strand NGS pipeline, Model-based Analysis for ChIP-Seq (MACS, version 2.0, Zhang et al., 2008)) with a p-value cutoff set to 1.0E-05, quality threshold >=30, 99% match to the sequence, and all duplicates removed (Fig. 3C). We also found coincident peak detection using ENCODE datasets from NRF-1 ChIP-seq with genomic DNA from immortalized cell lines (K562, HepG2, CH12.LX, GM 12878, and HeLa-S3; data not shown). Note that the detected peak in H1-hESC is identical to that predicted by Figure 2 and within the wildtype oligonucleotide sequence that bound nuclear extracts from rat primary neurons (Fig. 3).

### Overexpression of NRF1 Induces GABRB1 Promoter Activity in Transfected Primary Cortical Neurons

To evaluate whether there is a functional consequence to NRF-1 binding to its consensus site in *GABRB1-p*, primary cortical neurons were transfected with the *GABRB1p*-*luciferase* construct containing the 541 bp 5’ flanking region upstream of the human β1 subunit gene (Russek et al. 2000). We chose this approach to study functional relevance of the NRF-1 site to *GABRB1* transcription in neurons because NRF-1’s influence on the genome is difficult to detect by siRNA knockdown due to its robust expression at baseline and protein stability (Baar *et al*. 2003, Scarpulla 2006, Ramachandran et al. 2008).

As the expression of the NRF1:VP16 fusion protein has been shown to induce the promoter activity of NRF-1 responsive genes in cell lines (Ramachandran et al. 2008, Gonen & Assaraf 2010), we transfected primary cortical neurons with NRF1:VP16 along with the *GABRB1p-luciferase* reporter and found a marked increase (∽70%, fold change: 1.671 ± 0.404, n=5, *p < 0.05) above baseline (when compared to co-transfection with empty vector control, 1.00 ± 0.225, n=5) (Fig. 4A). Mutations were introduced into *GABRB1-p* using site-directed mutagenesis (based on the loss of specific binding of NRF-1 as identified in EMSA (see Fig. 3, lane 8)). As can be seen in Figure 4A for *mGABRB1-p*, with and without NRF1:VP16 overexpression, mutation of the NRF-1 regulatory element in *GABRB1-p* reduces basal activity to around 30% (fold change: 0.314 ± 0.067, n=5, *p < 0.05) of wild type. Overexpression of NRF-1:VP16 has no effect on *mGABRB1-p* (0.358 ± 0.057, n=5, ns) showing that increased *GABRB1* promoter activity directed by NRF-1 is sequence specific; and, moreover, that NRF-1 may be an important positive regulator of β1 subunit expression in developing neurons, especially interesting because β1 is found in the germinal zones and associated with pre-migrating neurons (Ma and Barker, 1995). Furthermore, increased mitochondrial biogenesis has also been associated with neuronal differentiation (Vayssiere *et al*. 1992, Cheng *et al*. 2010).

**Figure 4.**
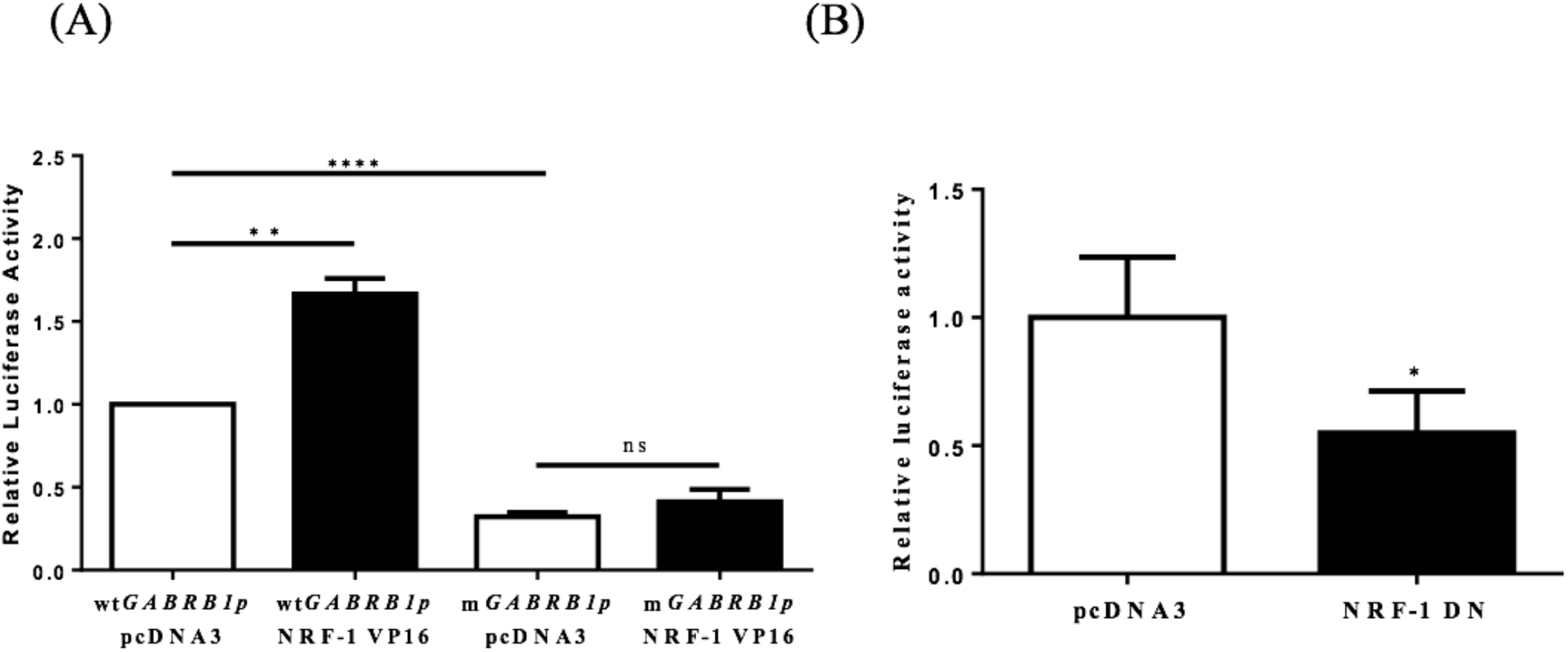
Evidence for the regulation of the GABAR °1 promoter by NRF-1. (A) Primary cortical neurons were co-transfected with 2 µg of wild type *GABRB1p* (*wtGABRB1p*) or the NRF-1 binding site mutant (*mGABRB1p*) and 1 µg of empty vector pcDNA3 or the NRF-1:VP16 fusion construct. Cells were assayed for luciferase activity 24 hours after transfection. Data represent the average ± SEM (n = 5 independent transfections) of luciferase activity relative to wild type *GABRB1p* in the absence of NRF-1:VP16. “*”and “ns” represent presence or absence respectively of significance based on p < 0.05 according to Student’s t-test. (B) Primary cortical neurons were co-transfected with either pcDNA3 or the dominant negative variant of NRF-1 (NRF-1 DN) and *GABRB1p* reporter (2 µg). Twenty-four hours after transfection, cells were assayed for luciferase activity. Data represent the average ± SEM of n = 6 independent transfections, normalized to *wtGABRB1p* and pcDNA3. *, p < 0.05, Student’s t-test. This dataset was originally published in the thesis of Li, Zhuting (2015) Activity-dependent gene regulation in neurons: energy coupling and a novel biosensor [dissertation]. [Boston (MA)]: Boston University.

### Inhibition of NRF-1 Function in Neurons

To evaluate the specific effect of endogenous NRF-1 on *GABRB1* transcription, a dominant negative form of NRF-1 was utilized that contains the DNA binding domain but lacks the NRF-1 trans-activation domain (Gugneja *et al*. 1996). Co-expression of this dominant negative NRF-1 represses *GABRB1* promoter activity by 45% (fold change: 0.549 ± 0.164, n=6, *p < 0.05, Fig. 4B) compared to empty-vector control (1.000 ± 0.235, n=6). Most importantly, overexpression of dominant negative NRF-1 blocks the activity dependent increase of endogenous *Gabrb1* mRNA levels in response to KCl treatment (Fig. 5).

When taken together with the fact that the NRF-1 element in the β1 subunit gene is completely conserved across species (Fig. 2), and that there is a mutation-induced loss of binding (Fig. 3) and function (Fig. 4), our results strongly suggest that NRF-1 is an essential feature of β1 subunit expression in neurons and that it couples transcription to the activity pattern of individual cells.

## DISCUSSION

We now report that the GABAR β1 subunit gene (*GABRB1*/*Gabrb1*) is regulated by NRF-1, a crucial transcription factor involved in oxidative phosphorylation and mitochondrial biogenesis. While it is believed that NRF-1 coordinates synaptic activity and energy metabolism by regulating excitatory neurotransmission via genes that code for subunits of the N-methyl-D-aspartate (NMDA) receptor (Dhar & Wong-Riley 2009, Dhar *et al*. 2009, Dhar & Wong-Riley 2011), it is clear that the this regulatory program is more complex than originally expected given our observation that elements of inhibitory neurotransmission may be coordinately regulated with excitation. This possibility is especially important as a variety of brain disorders present with a decrease in GABAR β1 subunit levels, including epilepsy where there is also aberrant hyperactivity.

Previously, our laboratory mapped the 5’ flanking region of the human β1 subunit promoter. Within this TATA-less promoter, we identified the major transcriptional start site (TSS) and described an initiator element (Inr) that senses the presence of prolonged GABA to mediate the autologous downregulation of β1 subunit expression (Russek et al. 2000). Our recent studies have discovered that such decreases in β1 subunit RNA levels may reflect a change in the chromatin state as mediated by PhF1b, a polycomb-like protein (Saha et al. 2013). In our present work, we have found a conserved canonical NRF-1 binding element (Fig. 2) that interacts with NRF-1 *in vitro* as verified by mobility shift assays (Fig. 3). Interestingly, our results in primary rat neurons are consistent with a peak of NRF-1 binding over the core promoter of *GABRB1* in human embryonic stem cells, as displayed in Figure 3C, using our bioinformatic analysis of ENCODE project datasets (Wang *et al*. 2012, Wang *et al*. 2013, Gerstein *et al*. 2012) with the Strand NGS pipeline (MACS V2 peak detection). We identified the same peak of binding in additional NRF-1 ChIP-seq ENCODE datasets from immortalized cell lines. Interestingly, we did not detect any additional NRF-1 peaks on the genes that code for other β subunit genes, suggesting that NRF-1 regulation may be unique to β1.

It is thought that NRF-1 binds as a homodimer to the consensus binding sequence (T/C)GCGCA(C/T)GCGC(A/G), making contact with DNA at the guanine nucleotides (Virbasius *et al*. 1993). This model is supported by the results of our mutational studies which show that a single mutation of G>C eliminates the ability of a cold double stranded oligonucleotide to compete for complex formation as assayed by mobility shift. The location of the NRF-1 element in *GABRB1* centers at −12 relative to the major TTS in neocortical neurons. The GC-rich NRF-1 binding motif is often associated with TATA-less promoters and found within 100 bp DNA regions around transcriptional start sites in the human genome (Virbasius et al. 1993, Xi *et al*. 2007). The proximity of the NRF-1 element in *GABRB1* to the Inr that binds polycomb-like proteins associated with chromatin remodeling and DNA methylation (Vire *et al*. 2006) may underlie its major role in controlling basal levels of β1 subunit mRNAs in neurons. Whether *GABRB1* is epigenetically regulated *in vivo* remains to be determined and could be a feature of why its transcription decreases in disease, especially since NRF-1 binding is blocked by DNA methylation (Gebhard *et al*. 2010).

Using the sensitivity of the luciferase reporter system, we have overexpressed NRF-1:VP16 in living neurons and shown that it indeed regulates the *GABRB1p-luciferase* reporter construct and that such regulation is lost upon mutation of the *GABRB1* NRF-1 regulatory element (Fig. 4A) and upon competition for endogenous NRF-1 binding to the promoter by overexpression of a dominant negative NRF-1 expression construct (Fig. 4B). We have also shown that the same mutation in the NRF-1 regulatory element of *GABRB1p* removes binding of endogenous NRF-1 to neuronal extracts in a mobility shift assay, as seen in Figure 3. Finally, and perhaps most importantly, we have shown that overexpression of dominant negative NRF-1 protein blocks the activity dependent increase in endogenous *Gabrb1* mRNA levels identifying a key molecular determinant of β1 subunit gene expression within cells (Fig. 5).

**Figure 5.**
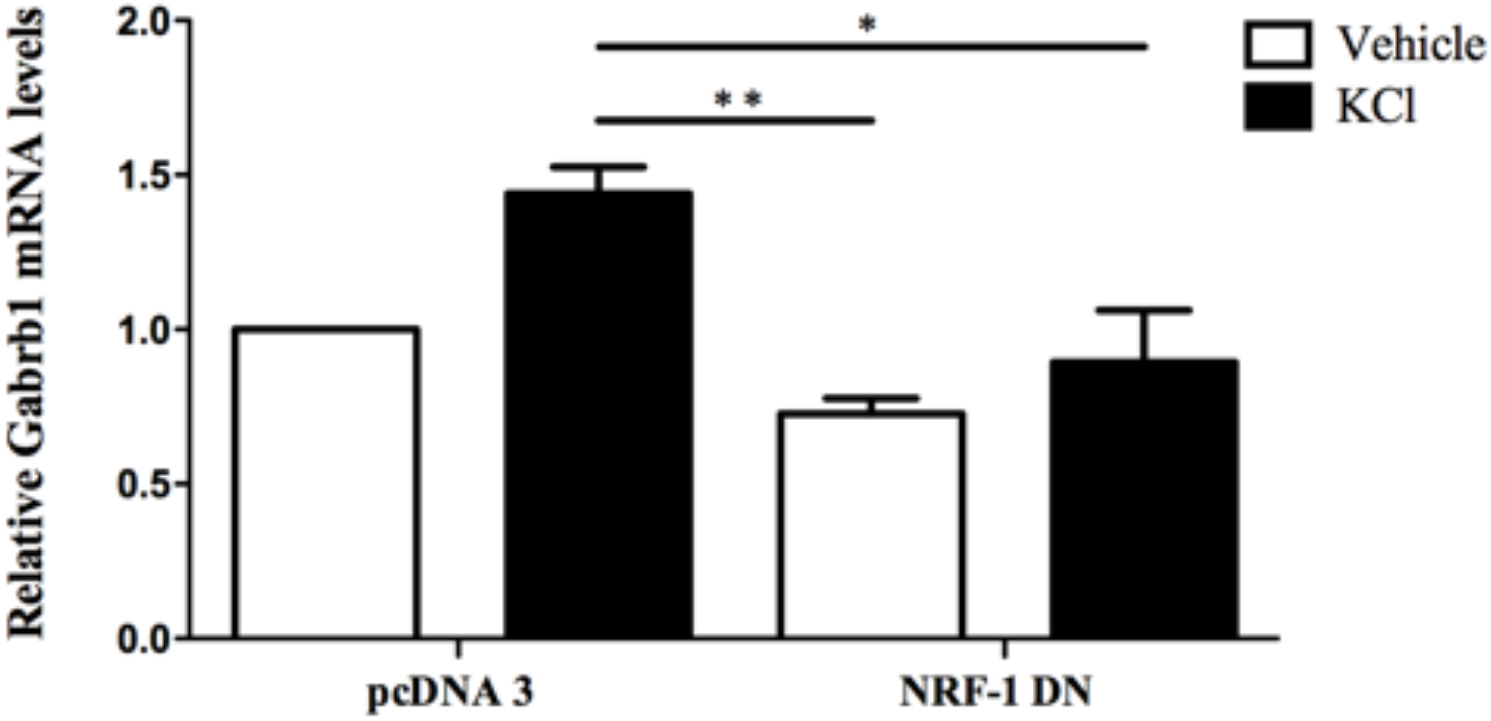
Overexpression of dominant negative NRF-1 attenuates the increase in β1 subunit mRNA levels in response to neuronal stimulation of primary cortical neurons. Primary cortical neurons were transfected with empty vector (pcDNA3) or dominant negative NRF-1 (NRF-1 DN) (using Nucleofection™) and plated in 6-well plates. DIV7 cells were treated with either vehicle or 20 mM KCl for 6 hrs. Total mRNA was isolated from cells and quantified by TaqMan qRT-PCR. *Gabrb1* mRNA expression was normalized to Cyclophilin A mRNA levels and is presented relative to its levels in pcDNA3 transfected neurons that were treated with vehicle (expressed as 1). Data represent the average ± SEM of n = 3 independent neuronal cultures. *, p < 0.05,**, p < 0.01, One-way ANOVA with Tukeys post hoc analysis.

Our results are consistent with previous studies from the Russek laboratory, using the same wild type *GABRB1p-luciferase* reporter construct, where promoter truncation and/or deletion that removes the Inr and at the same time disrupts the element for NRF-1 results in a 75–90% decrease in luciferase gene transcription (Russek et al. 2000) (Fig. 4). Given that GABAR blockade by bicuculine has also been shown to drive NRF-1 dependent transcription (Delgado & Owens 2012) and that bicuculine reverses GABA-induced downregulation of β1 mRNA levels (Russek et al. 2000), presumably through PhF1b binding to the Inr, our new results suggest that the NRF-1 responsive element and Inr may act synergistically to regulate β1 subunit levels as neurons adapt to changes in their activity state.

The direct regulation of NRF-1 in *GABRB1*/*Gabrb1* gene expression in the brain may also have implications in the initiation of sleep. The use of fragrant dioxane derivatives that show a 6-fold preference for β1-containing GABARs (Sergeeva *et al*. 2010) suggest that the β1 subunit is required for the modulation of wakefulness that is mediated by the histaminergic neurons of the posterior hypothalamus (tubermamillary nucleus-TMN) (Yanovsky *et al*. 2012). Given that energy metabolism is sensitive to restoration during the sleep cycle and that NRF-1 levels rise with sleep deprivation (Nikonova *et al*. 2010), it is interesting that β1-containing GABARs are the major source of inhibitory control over sleep.

Although differential expression of α subunits in relationship to brain disorders has clearly been associated with their region-specific control over changes in tonic and phasic inhibition, it is only recently that the importance of differential β subunit expression to GABAR function has been noted. This selective property of GABAR function ascribed to the assembly of particular β subunits, however, has been limited to β2 and β3, with β1 present in only a limited population of receptors in the brain. However, of all β subunits, β1 has been most associated with both neurological and neuropsychiatric disorders. The reason for this functional relationship remains to be described and is an active area of investigation in our laboratories.

## Abbreviations

ChIP, Chromatin-immunoprecipitation; DIV, days *in vitro*; EMSA, Electrophoretic mobility shift assay; GABAR, GABA type A receptors; *GABRB1*, GABA receptor subtype A β1 subunit gene; HRP, horseradish peroxidase; Inr, Initiator element; NRF-1, Nuclear Respiratory Factor 1; PhF1, Polycomb-like protein; TSS, Transcriptional start site; VP16, herpes simplex virus virion protein 16.

## Acknowledgments

We thank Dr. Tod Gulick (Sanford-Burnham Medical Research Institute, Orlando, FL) for his helpful discussion and providing us with expression constructs that were invaluable for our experiments.

